# Molecular Typing with COI - DNA Barcode of mosquitoes with medical importance from rural area of La Pintada, Antioquia, Colombia

**DOI:** 10.1101/260505

**Authors:** Richard Hoyos-López

## Abstract

DNA barcode is a methodology that allows the identification of species using a short fragment of cytochrome oxidase I and library sequences stored in the barcode of life database (bold>), make up an alternative tool for mosquito identification in areas epidemiologically active for arboviruses, protozoa and bacteria. In our study, we collected 114 adult mosquitoes in a rural area in the municipality of La Pintada (Antioquia, Colombia), and were separate for genus and species using morphological keys. Two Legs were taken of specimens mounted, and these were used for DNA extraction, amplification of COI-Barcode through PCR and sequencing. 38 sequences were characterized of seven mosquito species and used in bold> for molecular identification, subsequent characterization of genetic distances intra/interspecies, and MOTUs grouping by neighbor-joining analyses. Seven MOTUs were separate corresponding to seven species identify by morphological keys. bold> was able to identify five species, and two were identified to the genre. The following medically important mosquitoes were recorded in the rural area from La Pintada *(Antioquia): Aedes aegypti, Anopheles triannulatus, Coquillettidia nigricans, Mansonia titillans, Ochlerotatus angustivitatus, Psorophora ferox and Psorophora (Grabhamia)* sp.

---

Mosquitoes (family Culicidae), comprise a monophyletic taxon and diverse group with 3,490 species recognized in temperate and tropical regions of the world (Harbach & Howard 2007). 150 speciemainly belonging to genera *Anopheles, Aedes* and *Culex* have been incriminated as vectors of pathogens that cause disease in human populations, these pathogens include arboviruses (mainly Togaviridae, Flaviviridae, Bunyaviridae), filarial worms (helminths) and protozoa (*Plasmodium* spp.) (Gubler 2002; Harbach 2007).

Despite their medical importance, the taxonomy of mosquitoes is far from complete and is a critical step for epidemiological studies, vector incrimination, natural infection rates with pathogens and control/prevention measures (Harbach & Kitching 1998: Reinert *et al.* 2004; Cywinska 2006). Traditionally, morphology-based taxonomy have been used for species identification of mosquitoes, but this procedure is time consuming and not always sufficient for identification to the species level; therefore, is necessary a multidisciplinary approach to taxonomy that includes morphological, molecular and data distribution (Krzywinski & Besansky 2003).

An alternative methodology was developed by Hebert et al. (2003), which involves the analysis of short genomic regions that can discriminate morphologically recognized animal species (DNA barcodes), suggesting that mitochondrial gene cytochrome c oxidase subunit 1(COI) can serve as a uniform target gene for a bio-identification system (Valentini *et al.* 2008). The ability of DNA barcodes to identify species reliably, quickly and cost-effectively has particular importance in medical entomology, where molecular approaches to diagnoses species are often of great benefit in the identification of all life stages, from eggs to adults (Cywinska 2006). The fast identification of mosquitoes with medical importance is an urgent topic for increase the taxonomic knowledge between pathogens and their invertebrate vector insects (Besansky *et al.* 2003).

DNA barcode is an alternative tool successfully working with other insect vectors with medical importance as Simuliidae (Rivera & Currie 2009), Phlebotomine sandflies (Azpurua *et al.* 2010; Hoyos *et al.* 2012a; Kumar *et al.* 2012, Contreras *et al.* 2014) and Tabanidae (Cywinska *et al.* 2010), achieving the characterization and fast identification of species in epidemiologically active zones for diseases transmitted for those insects. In mosquitoes, there is a significant progress in the molecular typification of medically important species using this approach in some countries as Canada (Cywinska *et al.* 2006), India (Kumar *et al.* 2007), Pakistan (Ashfaq *et al.* 2014), Argentina (Díaz-Nieto *et al.* 2013), Japan (Taira *et al.* 2012), Ecuador (Linton *et al.* 2013) and China (Wang *et al.* 2012). DNA barcode also been used in studies about mosquito species complex (Kumar *et al.* 2013), identification of potential vector of arboviruses (Golding *et al.* 2012; Hoyos et al. 2015a) and integrative taxonomy (Ruiz *et al.* 2010; Ruiz *et al.* 2012; Laurito *et al.* 2013).

Colombia has large areas of disease transmission involving mosquitoes; among them is malaria, dengue (DENV), yellow fever (YFV), Venezuelan equine encephalitis (VEEV) and other arboviruses of low circulation (Tinker & Olano 1993; Groot *et al.* 1996; Rodríguez *et al.* 1995; Rivas *et al.* 1995; Ferro *et al.* 2003; Mattar *et al.* 2005; Montoya-Lerma *et al.* 2011; Hoyos et al. 2015a; Hoyos et al. 2015b; Hoyos et al. 2015c; Hoyos et al. 2016). The Antioquia department is a geographic area with a significant diversity of ecosystems, reservoirs and natural hosts, that makes possible the emerging and re-emerging diseases and epidemic outbreak event; in this context, few researches have been realized studies about diversity of mosquito species in zones with ecological characteristic that allow natural circulation of emerging and re-emerging pathogens (Parra 2012; Hoyos *et al.* 2012b). In our study, we used DNA barcode methodology for molecular typing for mosquito species of medical importance present in the rural area on the municipality of La Pintada (Antioquia).

## MATERIALS AND METHODS

Mosquito collection. Mosquitoes used in this study were collected between February - April in 2012 in rural area from La Pintada (5°44’25.63” N, 75°36’20.18” W), department of Antioquia, Colombia. Adults were collected using CDC - light traps and manual aspirators close to rural forest patch. The traps were placed before sunset and insects were collected the following morning (6:00 p.m. to 6:00 a.m). Collected mosquitoes were stored in cryovials, placed in a liquid nitrogen tank and transported to the laboratory of Biology and Insect Systematics from National University of Colombia, Medellín. Specimens were separated considering genus, external morphological characteristics, date/site collection, for facilitate the morphological identification with pictorial keys (Gabaldon *et al.* 1946; Lane 1953; Cova-García *et al.* 1966; Forattini 2002) and molecular procedures.

Molecular protocols. Two legs were taken from selected mosquitoes identified and used for DNA extraction with commercially available DNeasy Blood & Tissue kit (Qiagen, Maryland). Amplification of the ~658 nt fragment of DNA barcode region from mitochondrial Cytochrome oxidase I gene was achieved using the primer pair LCO-1490/HCO-2198 (Folmer *et al*. 1994; Hebert *et al.* 2003. Each PCR-mix contained 1x NH4SO4 buffer, 1 mM each DNTP, 5 mM of MgCh, 0.5 uM each primer, 0,4 U of taq polymerase (Bioline, Maryland) and 4 uL of DNA template and was made up to a total volume of 50 uL using water for molecular biology quality. The PCR thermocycler parameters included: a single cycle at 94°C for 10 min, followed by 35 cycles of 95°C for 60 s, 50°C for 60 s and 72°C for 60 s, respectively terminating with a 72°C for 5 min of final extension step and 4°C hold. PCR products were visualized on 1% agarose gels, containing GELSTAR^®^ (Lonza, Rockland) diluted 1/50 using a Dark Reader lector (IMGEN, Alexandria). PCR products were sequenced using same primers LCO/HCO in Macrogen sequencing service (Seul, Korea).

Data analyses. Sequences were edited manually using Bioeditv7.2.0 (http://www.mbio.ncsu.edu/BioEdit/bioedit.htm) and aligned in ClustalW (Larkin *et al.* 2007). Genetic distances was calculated in MEGAv6.0 (Tamura *et al.* 2013) using the kimura 2-parameter distance model (K2P) (Kimura 1980), and molecular operational taxonomic units (MOTUs) was evaluated using clusters on dendrogram and genetic distances. In this sense, a neighbor-joining tree (NJ) (Saitou & Nei 1987) was generated using K2P and bootstrap (1,000 replicates) (Felsenstein 1982). For some species with more than four sequences were calculated estimates of genetic diversity as diversity haplotype and polymorphic sites, using DNAspv5.0 software (Librado & Rozas 2009).

## RESULTS

A total of 114 mosquitoes were collected in the rural area from municipality of La Pintada (Antioquia), and seven species mosquitoes were identified: *Ochlerotatus (Ochlerotatus) angustivitatus* (n = 52) (Dyar & Knab 1907), *Mansonia (Mansonia) titillans* (n = 13)(Walker 1848), *Aedes (Stegomyia) aegypti* (n = 18) (Linnaeus 1762), *Psorophora (Janthinosoma) ferox* (n = 14)(Von Humboldt 1819), *Coquillettidia (Rhynchotaenia) nigricans* (n = 9) (Coquillett 1904), *Anopheles (Nyssorhynchus) triannulatus* (n = 1) (Neiva & Pinto 1922) and *Psorophora (Grabhamia)* sp (n = 7).

38 DNA barcode sequences (19 haplotypes) belonging to seven species representing six genera were obtained and compared to those available in bold> database for verify taxonomic identifications; four species were confirmed, two only were possible determine genera because species was not released in the database, and one not show homology with any other mosquitoes sequence (Table I). Intra-specific genetic distances and haplotypes in seven species were low and within ranges previously reported as corresponding like inside species (0 - 0.01). Interspecies genetic distance it were consistent and in the range between species from different genera (Table II). However, genetic variability of mosquitoes was low in haplotypes also showing a low number of polymorphic sites. The separation of mosquito species by genetic distances was confirmed through the neighbor-joining dendrogram and MOTU’s grouped (Fig. 1).

**Table I.**
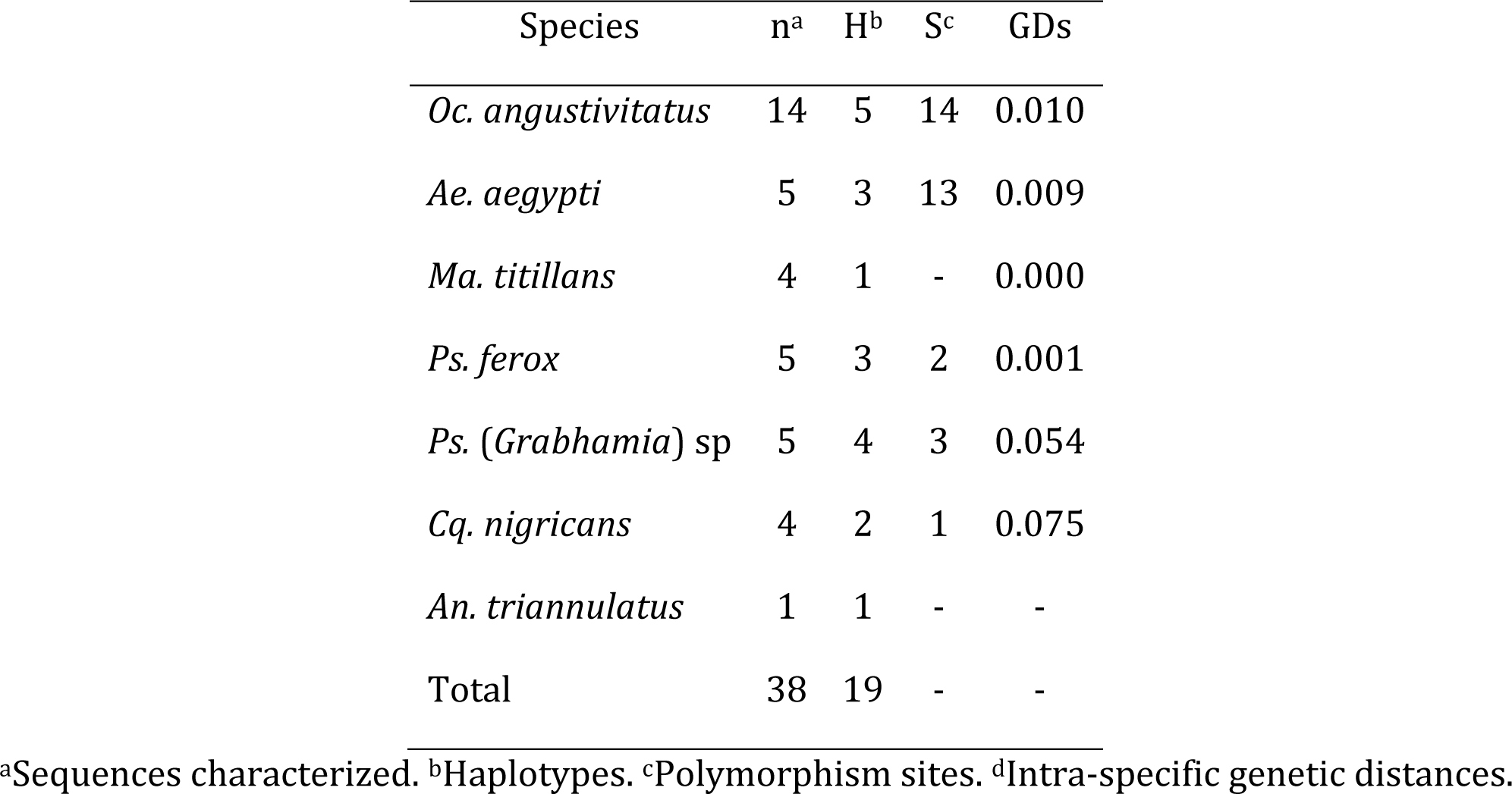
Diversity of DNA barcode sequences for seven species of mosquitoes collected in rural area from La Pintada.

**Table II.** Genetic distances between DNA barcode sequences belonging to seven mosquito species identified in rural are from La Pintada (Antioquia).

|  | 1 | 2 | 3 | 4 | 5 | 6 | 7 |
| --- | --- | --- | --- | --- | --- | --- | --- |
| 1. *Ae. aegypti* |  | 0,017 | 0,016 | 0,016 | 0,015 | 0,015 | 0,017 |
| 2. *An. triannulatus* | **0,153** |  | 0,016 | 0,016 | 0,016 | 0,016 | 0,016 |
| 3. *Cq. nigricans* | **0,172** | **0,166** |  | 0,014 | 0,014 | 0,015 | 0,016 |
| 4. *Ma. titillans* | **0,146** | **0,152** | **0,135** |  | 0,016 | 0,017 | 0,018 |
| 5. *Ps (Grabhamia) sp* | **0,142** | **0,150** | **0,156** | **0,161** |  | 0,012 | 0,013 |
| 6. *Oc. angustitativatus* | **0,136** | **0,141** | **0,164** | **0,153** | **0,106** |  | 0,014 |
| 7. *Ps. ferox* | **0,137** | **0,133** | **0,173** | **0,176** | **0,108** | **0,110** |  |

Numbers in boldface indicate interspecific genetic distances and the others values show standard deviations.

**Figure 1.**
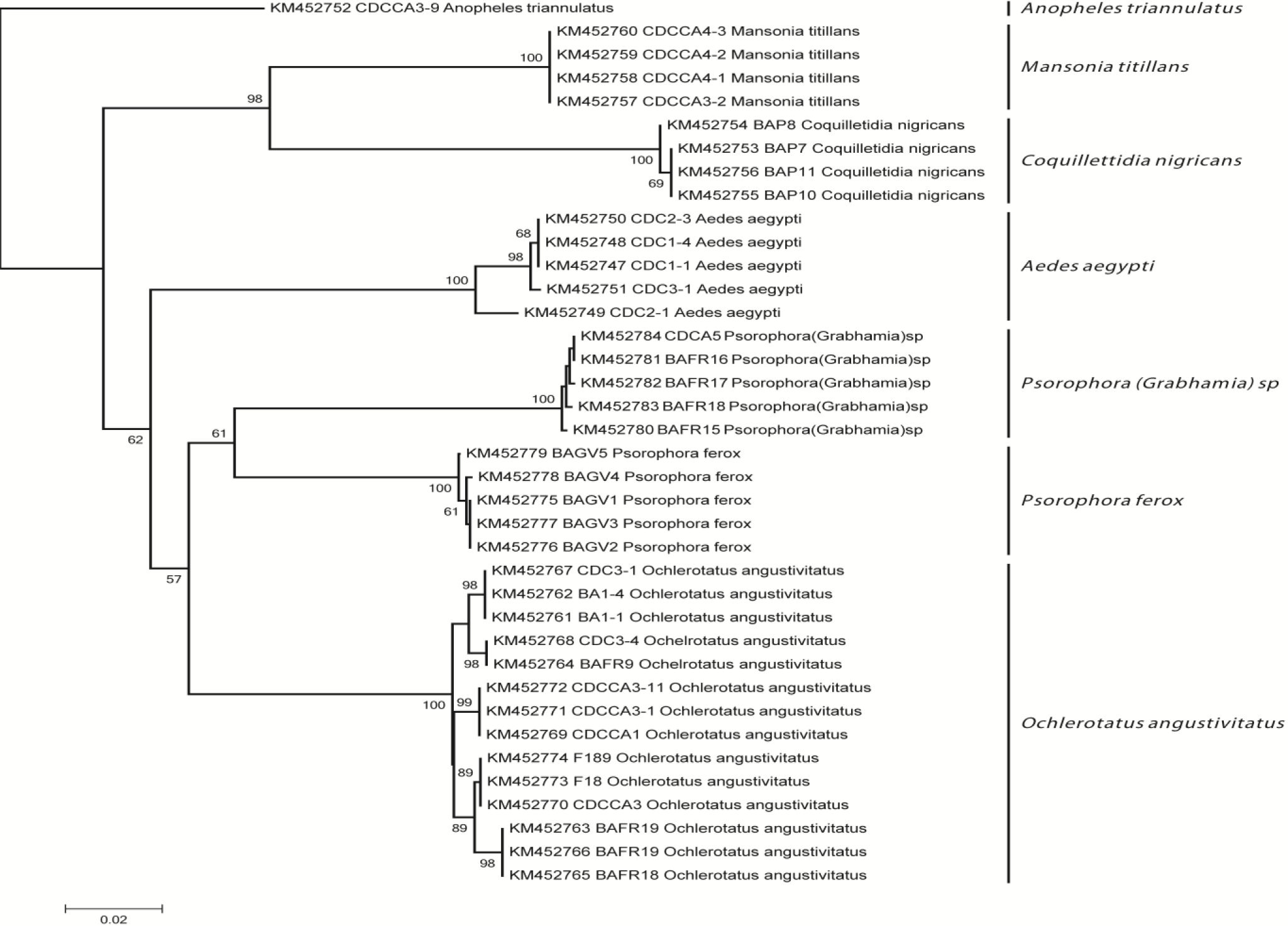
Neighbor-joining dendrogram using DNA barcode sequences (Cytochrome oxidase I) (Kimura two parameter genetic distances, Bootstrap = 1000 replicates). Percentage of replicate trees in which the associated taxa clustered together in the bootstrap test is show next to the branch (only >50%). The final dataset comprised 574 nucleotides, including all codon positions. Every sequence shows his number access Genbank before mosquito code and species identification.

*Ps.* (*Grabhamia*) sp, despite being separate with DNA barcode sequences from *Ps. ferox;* the females collected could not be identified morphologically to species, and males not were collected for taxonomic verification.

## DISCUSSION

Correct identification of the insect vector is the one most important factors in the study of the arboviruses and protozoa diseases (Cook *et al.* 2005; Besansky *et al.* 2003), and the precise identification of the target species has direct medical and practical implications, particularly in developing vector control strategies (Dhananjeyan *et al.* 2010; Zamora et al. 2015).

Taxonomy mosquito has been achieved mostly using morphological characteristics, cyto-genetics and iso-enzymes markers; recently, the molecular approaches have signficants improvement in the accuracy of species identification using DNA barcodes (Cywinska *et al.* 2006; Kumar *et al.* 2007; Ruiz *et al.* 2010; Golding *et al.* 2012; Ruiz *et al.* 2012; Wang *et al.* 2012). The Barcode - COI region in mosquitoes is characterized by a high rate of evolution (transitions/transversions) (Cywinska *et al.* 2006) allowing the estimation of biodiversity based in molecular operational taxonomic units (MOTUs) (Blaxter 2004).

In our study, seven MOTUs were separate by genetic distances (K2P) and tree building criteria with neighbor-joining dendrogram. Genetic distances showed similar values to reported within (0 - 0,010) and between species (0,108 - 0,176) in other studies (Cywinska *et al.* 2006; Kumar *et al.* 2007; Wang *et al.* 2012). Genetic diversity of DNA barcodes was low in mosquitoes collected; these results probably evidenced selective process that decreases haplotype diversity in natural populations by urban aspersion of insecticides and/or pesticides for rural crops and fragmentation of natural habitats (Ocampo & Wesson 2004; Yanoviak *et al.* 2006; Keesing *et al.* 2010).

The six species identified have a great relevance in epidemiology and medical entomology, because they have been founded infected with human pathogens as arboviruses and protozoa in Colombia and nearby countries:

- *Ae. aegypti* is responsible for the transmission of Dengue virus, the most important arboviruses in Colombia (Tinker & Olano 1993), and was vector of Yellow Fever virus in urban zones after his eradication (Rodriguez *et al.* 1996). The blood feeding pattern is anthropophilic, and is well adaptive to urban areas (Harrington *et al.* 2001).
- *Ps. ferox*, females of this species frequently carry eggs of *Dermatobia* in eastern of Colombia and have also been found infested with *Dermatobia* eggs in Panama (Capenter & LaCasse 1955). Groot *et al.* (1996), determines natural infection of mosquitoes from this species for Venezuelan encephalitis equine virus, Ilheus, Mayaro and Una virus in San Vicente de Chucurí (Santander); Wyeomyia virus was detected too in pooles of *Ps. ferox* from middle Magdalena valley. In other countries, *Ps. ferox* has been found with Jamestown Canyon virus (California, EEUU) (Andreadis *et al.* 2008), Venezuelan equine encephalitis virus in Alabama (EEUU) (Chamberlain *et al.* 1956), Mayaro (Brazil) (Muñoz & Navarro 2012), Rocio virus (Brazil) (Souza *et al.* 1981) and West Nile virus (EEUU) (Kulasekera *et al.* 2001; Andreadis *et al.* 2004).
- *Ma. titillans*, their larvae prefer small lakes, ponds, rivers with little stream and swamps, associate with floating plants as water lettuce - *Pistia stratiotes* - (Lounibos & Linley 2008). Blood feeding patterns showing are broad spectrum with a tendency to mammals and birds (Edman & Kale 1971). Parra *et al.* (2012) recorded this species in forest fragments from Uraba (Antioquia) closed to human habitats and Groot *et al.* (1996), found insects infected with Venezuelan encephalitis equine virus in Magangué (Bolivar) in 1970-1971. In alligator farms from Florida (EEUU) had been register with West Nile virus (Unlu *et al.* 2010).
- *An. triannulatus* is a species complex evidenced with variations in male genitalia, eggs, larvae and molecular markers (Rosa-Freitas *et* al. 1998; Silva do Nascimento *et al.* 2002; Silva do Nascimento *et al.* 2006; Rosero *et al.* 2012; Moreno *et al.* 2013), these mosquito species is implied in transmission of *Plasmodium vivax* in Brazil (Benarroch 1931; Gabaldon *et al.* 1946; Oliveira-Ferreira *et al.* 1990; Tadei *et al.* 2000), Perú (Aramburú *et al.* 1999) and recently in the locality of La Capilla (El Bagre, Antioquia) from Colombia (Naranjo *et al.* 2013).
- *Cq. nigricans.* The habits of this mosquito are similar to *Mansonia* species and founded naturally infected with the genotype IE Caribbean/Gulf of Venezuelan equine encephalitis virus in Mexico (Adams *et al*. 2012).
- *Oc. angustivitatus.* Females are founded in environments sylvatic, peridomiciliary and inside houses, with bite-tendency to equines and domestic animals (Parra et al 2012). In Panamá had been recorded with Ilheus virus and Venezuelan equine encephalitis virus in Colombia (Forattini 2002).

DNA barcode is a successful tool for rapid identification of mosquitoes, biodiversity estimation, epidemiological studies on unexplored zones, and ecology of emerging diseases transmitted by mosquito vectors, however, is indispensable use different approximations for the analysis of sequences and not over/under-estimate molecular diversity (Meier *et al.* 2006; Blaxter 2004; Valentini *et* al. 2008). In our study, we achieved identify several species fundamentally mosquitos associate to rural areas, about these species, there is enough sequences in bold> for a clear identification using different kind analysis, but sylvatic/forest species are more difficult as noted in rural areas from La Pintada, in part is due to lack of sequences from mosquitoes identified belonging to these unexplored ecosystems, however, DNA barcode allowed molecular separation, as be observed in *Ps. (Grabhamia)* sp.

Some species founded in La Pintada (Antioquia) as *Ps. ferox* and *Ma. titillans* are species with long dispersal ability from fragment forest to open areas (Mendez *et al.* 2001), allowing dispersion of arboviruses to susceptible hosts (adjacent human population and domestic animals) and vectors adapted to artificial ecosystems (farms, field crops), as *Ae. aegypti* populations could be origin to epizootia or epidemic break in humans (Tanaka *et al.* 1979; Tinker & Olano 1993; Oda *et al.* 1999; Pecor *et al.* 2002; Hoyos et al. 2015d). The ecological context present in La Pintada (Antioquia, Colombia), make this locality susceptible to emerging and re-emerging pathogens, due to changes in community of mosquitoes in rural and urban environments (Yanoviak *et al.* 2006, Keesing *et al.* 2010).

Is recommended begin more research in bite-behavior rates, detection of emerging and re-emerging pathogens (alphavirus and flavivirus groups), and natural habitats (phytotelmata), for identify human epidemiological risk in rural populations and establish future measures of control and prevention for diseases.

## ACKNOWLEDGEMENTS

To Diego Arias and Diego Puerta for his assistance in entomological surveys and laboratory work.

